# Comparison of the neuraminidase antigenicity in recently circulating influenza A and vaccine viruses

**DOI:** 10.1101/2021.05.21.445157

**Authors:** Jin Gao, Xing Li, Hongquan Wan, Zhiping Ye, Robert Daniels

## Abstract

Neuraminidase (NA or N) antigens in circulating influenza viruses are not extensively evaluated for vaccine strain selection like hemagglutinin (HA or H) even though viral-based influenza vaccines include the recommended strain NA in varying amounts. As NA can also elicit a protective response, we assessed the antigenic similarity of the NAs from human H1N1 and H3N2 viruses that were prevalent between September 2019 to December 2020 to NAs from several recently recommended vaccine strains. To eliminate the dependence on isolates, the enzyme-linked lectin assay for analyzing NA antigenicity was performed with reverse genetic viruses carrying the same HA. Our results show that ferret antisera against NAs from the recommended H1N1 and H3N2 vaccine strains for the 2020-21 northern hemisphere influenza season recognize and inhibit the most prevalent circulating N1s and N2s, suggesting the NAs from the influenza A vaccine and circulating strains are antigenically similar. Comparisons of the recent N2s also revealed a bias in the reactivity of NA antisera from the egg and cell-based H3N2 vaccine strains due to a C-terminal substitution, indicating the C-terminus can influence N2 antigenicity and should receive consideration during the H3N2 strain selection.

## INTRODUCTION

The first licensed U.S. influenza vaccine was bivalent, consisting of an influenza A virus (IAV) and an influenza B virus (IBV) strain that were propagated in chicken eggs, isolated using red blood cells, and inactivated with formalin [1, 2]. During these early trials vaccinated individuals were reported to be more protected and to generate antibodies capable of inhibiting the receptor binding function of the hemagglutinin (HA or H) antigen, a property which eventually was established as a correlate of protection for influenza vaccines [3]. Around the time hemagglutination-inhibition (HAI) was accepted, several studies began to show that antibodies against the less abundant surface neuraminidase (NA or N) antigen can also limit the severity of influenza infections in animals and humans [4–6]. However, establishing a correlate of protection for NA and quantifying the content in vaccines have proven to be more difficult than for HA.

Inactivated viruses are still the most common form of influenza vaccines administered in the U.S. [7]. The vaccine viruses are now produced in cells, as well as eggs, and the purification and inactivation processes have improved [8–10]. Recently, most viral-based influenza vaccines are also quadrivalent, containing HA and NA antigens from two IAV and two IBV recommended strains [11]. The two IAV strains are from the H1N1 and H3N2 subtypes, and the two IBV strains are from the Yamagata and Victoria lineages. To account for changes in circulating influenza viruses, the vaccine strains are selected annually based on a combination of epidemiological, HA antigenicity and modelling data [12]. Efforts are then made to create high growth reassortant candidate vaccine viruses (CVVs) for cell and egg-based vaccines that possess a HA which is antigenically similar to the recommended strain HA.

Despite the continuous strain evaluations and updates, the efficacy of influenza vaccines remains less than ideal [13, 14], suggesting influenza vaccine improvements cannot be achieved by HA alone. Consequently, NA is receiving more consideration as a second influenza vaccine antigen, especially since NA antibodies can also provide cross protection against strains carrying HAs that are antigenically distinct from the vaccine [15–19]. However, NAs in vaccines are currently not selected or evaluated, but rather a consequence of the pairing with a recommended HA in an available isolate. As a result, cell and egg-based vaccine strains for the same influenza virus subtype or lineage can contain distinct NAs. Another major obstacle for evaluating NA is the general lack of antigenic data for NA in circulating strains.

Many recent studies have begun to use an immobilized enzyme-linked lectin assay (ELLA) to measure NA inhibition (NAI) antibody titres and to assess NA antigenicity [20–25]. ELLA was first developed to examine NA activity and inhibition using erythrocyte glycoproteins as a substrate and agglutination by the galactose binding lectin peanut agglutinin as a reporter [26, 27]. This assay has since been modified and the glycans from bovine fetuin are used for measuring viral NA activity in the absence and presence of serum containing NA antibodies [28–32]. An advantage of ELLA is that it can capture antibody-mediated interference of NA enzymatic activity [33, 34], which has been shown to correlate reasonably well with NA-mediated protection [21, 35]. However, in recent years the analysis of H3N2 isolates by ELLA has been very limited because of these strains grow poorly in eggs [36].

In this study we compared the antigenicity of NAs subtype 1 (N1) and 2 (N2) from recently circulating IAVs and vaccine strains using ferret antisera coupled with ELLA. To eliminate the dependence on field isolates, the analysis was performed using reverse genetic viruses that all carry the same HA with different NAs from circulating IAV strains. Our results show that the NAs from the recently recommended H1N1 and H3N2 vaccine strains are antigenically similar to the NAs from the predominant circulating IAV strains and that the C-terminus of N2 should be considered during the H3N2 strain selection as this is an antigenic determinant. The benefit of combining bioinformatics and reverse genetics with the ELLA for evaluating NA antigens in current and future vaccine strains is discussed.

## RESULTS

### Enzyme-linked lectin assay (ELLA) for evaluating the antigenicity of recent NAs

To evaluate NA antigenicity we performed a standard ELLA that utilizes the glycans on immobilized bovine fetuin as the NA substrate, the galactose-binding lectin peanut agglutinin as the reporter, and reverse genetic viruses grown in embryonated chicken eggs as the NA source (Fig. 1A and [31, 32, 37]. Although generating reverse genetic viruses eliminated the dependence on strain isolates and minimized the potential ELLA interference of antibodies against the inoculating strain HA [38, 39], the viruses were still likely to vary in NA content. Therefore, we first compared the ELLA results from two reassortant viruses (PR8^H1N1-BR18^ and WSN^H1N1-BR18^) that possess different amounts of the same HA and NA (Fig. 1B and [25]). Based on Coomassie stained gels (Fig. 1C), NA activities and hemagglutination unit (HAU) titres (Fig. 1D), the WSN^H1N1-BR18^ virus was estimated to contain ~2.5 times more NA and less HA than the PR8^H1N1-BR18^ virus. Supporting these properties, NA-mediated release of WSN^H1N1-BR18^ from fetuin was faster than PR8^H1N1-BR18^, whereas the HA binding affinity was slightly less (Fig. 1E). The ELLA captured the ~2-fold higher NA content in the WSN^H1N1-BR18^ virus (Fig. 1F, upper panel) and produced similar NAI titres for both viruses when it was coupled with ferret antisera against N1-BR18 (Fig. 1F, lower panel, and Table 1), confirming our experimental setup can account for changes in the viral NA content when the HAs are identical.

**Figure 1.**
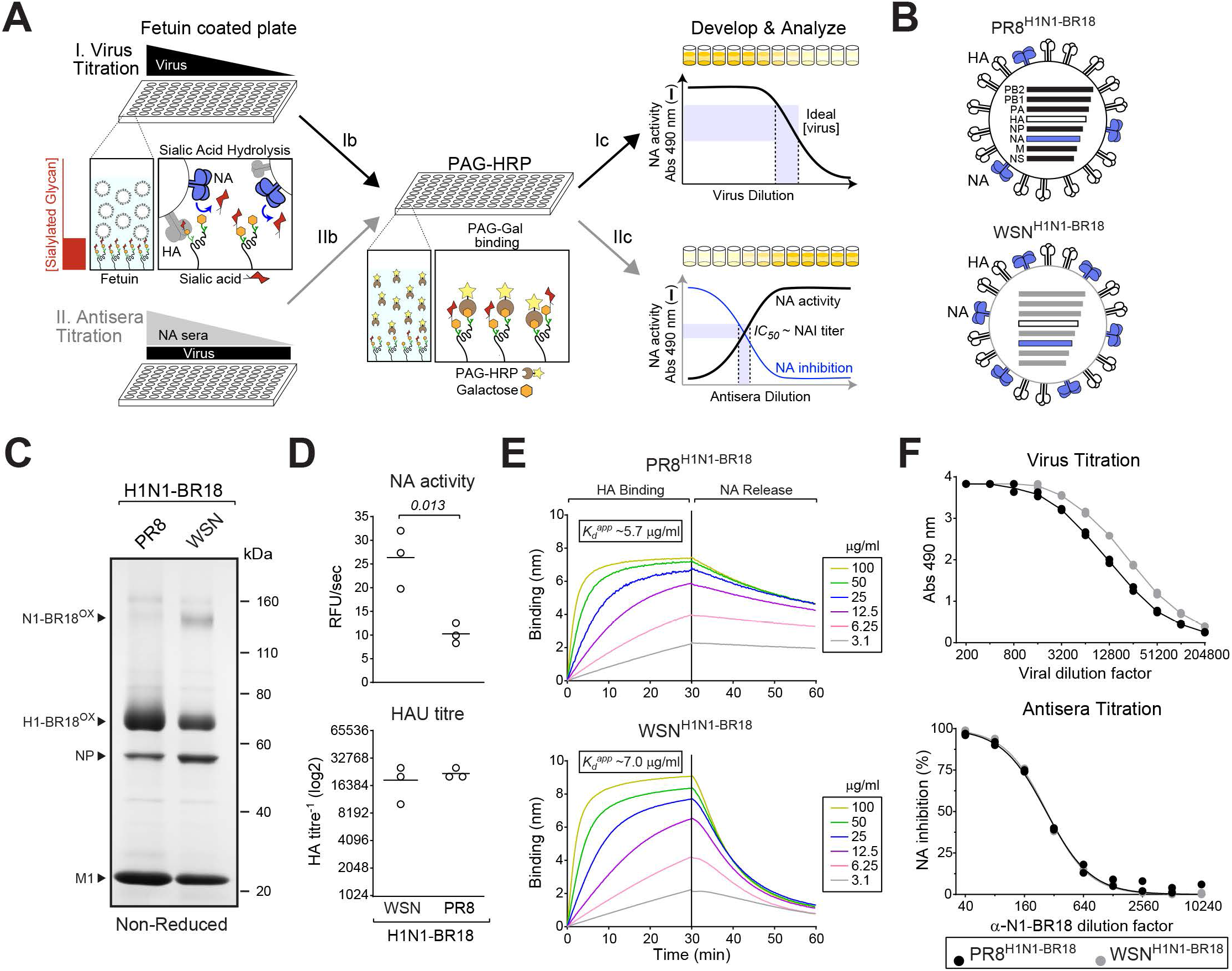
The enzyme-linked lectin assay accounts for variation in viral NA content. **A.** Illustration of the ELLA for evaluating NA. (I) Viruses are titrated using fetuin coated plates. (Ib) The NA exposed galactose residues on the fetuin glycans are bound by peanut agglutinin linked to HRP (PAG-HRP). (Ic) PAG-HRP binding is determined using the substrate OPD and measuring the Abs at 490 nm. (II) Antisera are then titrated with a fixed virus dilution from the linear range, added to fetuin coated plates and processed similarly. (IIc) NA inhibition (NAI) titres are determined by plotting the NA activity data inversely and calculating the antisera 50% inhibitory concentration (*IC_50_*). **B.** Diagram of the PR8^H1N1-BR18^ and WSN^H1N1-BR18^ reassortant viruses. **C.** Non-reduced Coomassie stained gel containing ~5 μg of purified PR8^H1N1-BR18^ and WSN^H1N1-BR18^ viruses. Oxidized (OX) N1-BR18 and H1-BR18 are indicated with the viral proteins NP and M1. **D.** NA activities (top) and HAU titres (panel) are shown for 3 independently purified virus batches with protein concentrations of 1 mg/ml. Significant *P* values from an unpaired student T-test, 95% CI, are displayed. **E.** Binding of purified PR8^H1N1-BR18^ (upper panel) and WSN^H1N1-BR18^ (lower panel) virions to fetuin was measured by BLI. Curves were generated with equal densities of immobilized fetuin and the indicated purified virus protein concentrations. HA binding was measured in the presence of 10 μM zanamivir to inhibit NA activity and calculate the apparent dissociation constant (*K_d_^app^*) for fetuin. To follow NA-mediated release, biosensors with bound viruses were moved to a well without zanamivir. **F.** The ELLA virus titration is displayed for the PR8^H1N1-BR18^ and WSN^H1N1-BR18^ viruses together with the ferret α-N1-BR18 antisera titration data (lower panel) that was used to calculate the NAI titre. Data from each sample run in duplicate are displayed.

**Table 1.** Reciprocal NAI titres from the indicated reassortant viruses and antisera. NAI titres were determined by calculating the *IC_50_* of the indicated ferret antisera by the ELLA. Cells where the NA in the PR8 or WSN reassortant virus matches the sequence of the NA used to generate the ferret antisera are highlighted. All ferret antisera were produced via intranasal inoculation with PR8 reassortant viruses carrying the indicated NA and H6. Samples were run in duplicate and the NAI titre is the mean from the two runs. NAI titres that could not be determined were listed as less than the lowest sera dilution. *n.d*.- not done.

|  |  | Ferret Antisera |  |  |  |
| --- | --- | --- | --- | --- | --- |
| | | α-N1-BR18 | α-N1-GD19 | α-N2-HK19e | α-N2-HK19c |
| Viruses | WSN<sup>H1N1</sup>-BR18 | 270.5 | *n.d.* | *n.d.* | *n.d.* |
|  | PR8<sup>H1N1</sup>-BR18 | 266.9 | *n.d.* | *n.d.* | *n.d.* |
|  | PR8<sup>N1</sup>-GD19 | *n.d.* | 6098 | <10 | <10 |
|  | PR8<sup>N2</sup>-HK19e | *n.d.* | <10 | 130 | 41 |
|  | PR8<sup>N2</sup>-HK19c | *n.d.* | <10 | 124 | 304 |

For the 2020-21 northern hemisphere influenza season, the NA sequences in the recommended H1N1 IAV strains for egg (A/Guangdong-Maonan/SWL1536/2019) and cell-based (A/Hawaii/70/2019) vaccines are identical. However, the NAs in the recommended H3N2 strains for egg (A/Hong Kong/2671/2019) and cell-based (A/Hong Kong/45/2019) vaccines differed at two positions (45 and 469). Based on these observations, we generated ferret antisera against one N1 vaccine antigen (N1-GD19) and the N2 antigens from the egg (N2-HK19e) and cell-based (N2-HK19c) vaccine strains. Each antiserum was produced by intranasal inoculation with a PR8 reassortant virus containing a HA (H6) gene segment and one of the vaccine strain NA gene segments (Fig. 2A). The antiserum specificity was tested by the ELLA using PR8 reassortant viruses carrying the HA (H1) gene segment from PR8 and the vaccine strain NA gene segments. The N1-GD19 antiserum showed high reactivity against PR8^N1-GD19^, producing an NAI titre of ~6000 (equivalent to the *IC_50_*), and minimal reactivity against the viruses carrying the N2 gene segments (Fig. 2B and Table 1). The N2-HK19e and N2-HK19c antisera produced modest NAI titres of ~130 and ~300 for their respective matching strains and did not react with the virus carrying N1-GD19 (Fig. 2B and Table 1). The N2 antisera also exhibited some bias for the strains with the matching N2s (Table 1), suggesting residues 45 or 469 could be in a N2 epitope for NAI antibodies.

**Figure 2.**
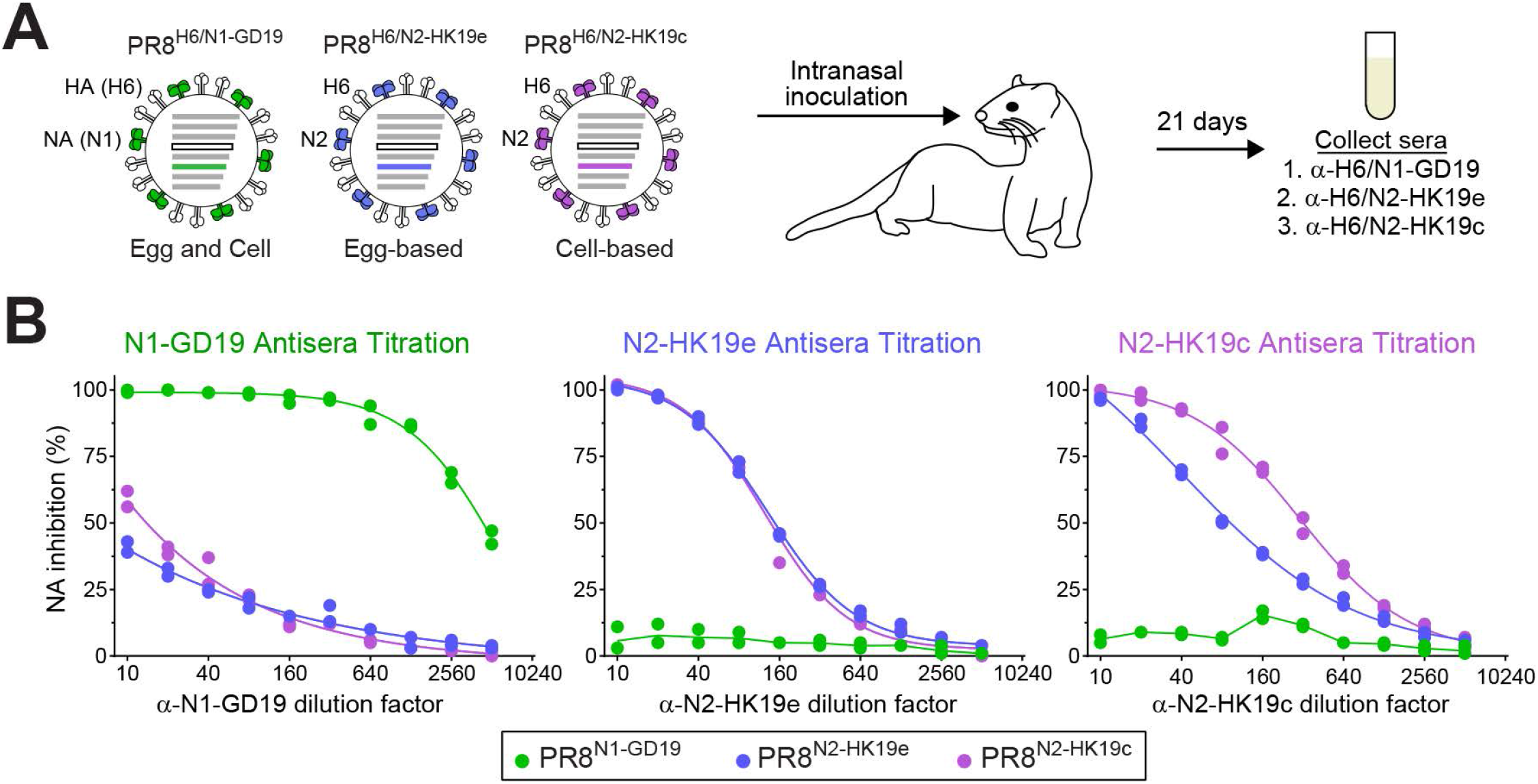
Characterization of ferret antisera against the NA vaccine antigens from the 2020-21 season. **A.** Diagram for generating ferret antisera against the NA antigens (N1-GD19, N2-HK19e, and N2-HK19c) from the WHO recommended egg and cell-based IAV vaccine strains for the 2020-21 northern hemisphere season. Ferret sera were collected 21 days post intranasal inoculation with an PR8 double-gene reassortant virus containing the NA gene from a recommended vaccine strain and a HA (H6) gene segment. **B.** Specificity and reactivity of each ferret antiserum was assessed by the ELLA using three PR8 single-gene reassortant viruses with each one containing a NA from the recommended vaccine strains. Data from each sample, run in duplicate, are displayed.

### Sequence and antigenic comparison of NAs from recent H1N1 and vaccine viruses

To identify potential antigenic changes in the NAs from recent H1N1 IAVs we compared the available NA sequences that were collected between September 1, 2019 to December 2020 to the N1-GD19 sequence. The frequency of amino acid changes in the recent N1 sequences was rather high at several positions in the stalk region, but it was much lower in the positions of the enzymatic head domain (Fig. 3A). With the exception of two positions in the stalk, the amino acid differences were mainly associated with a change to a specific amino acid (Fig. 3B). Mapping the more prevalent substitution positions on the N1 head domain structure revealed that most are surface exposed. One position (467) is located on the top of the tetramer, another (222) is near the active site, and several are located on the side and the bottom of the tetramer where the head is linked to the stalk (Fig. 3C). The surface exposure, and the proximity of at least one position to the NA active site, indicated that these common N1 substitutions could result in antigenic changes that are detected by the ELLA.

**Figure 3.**
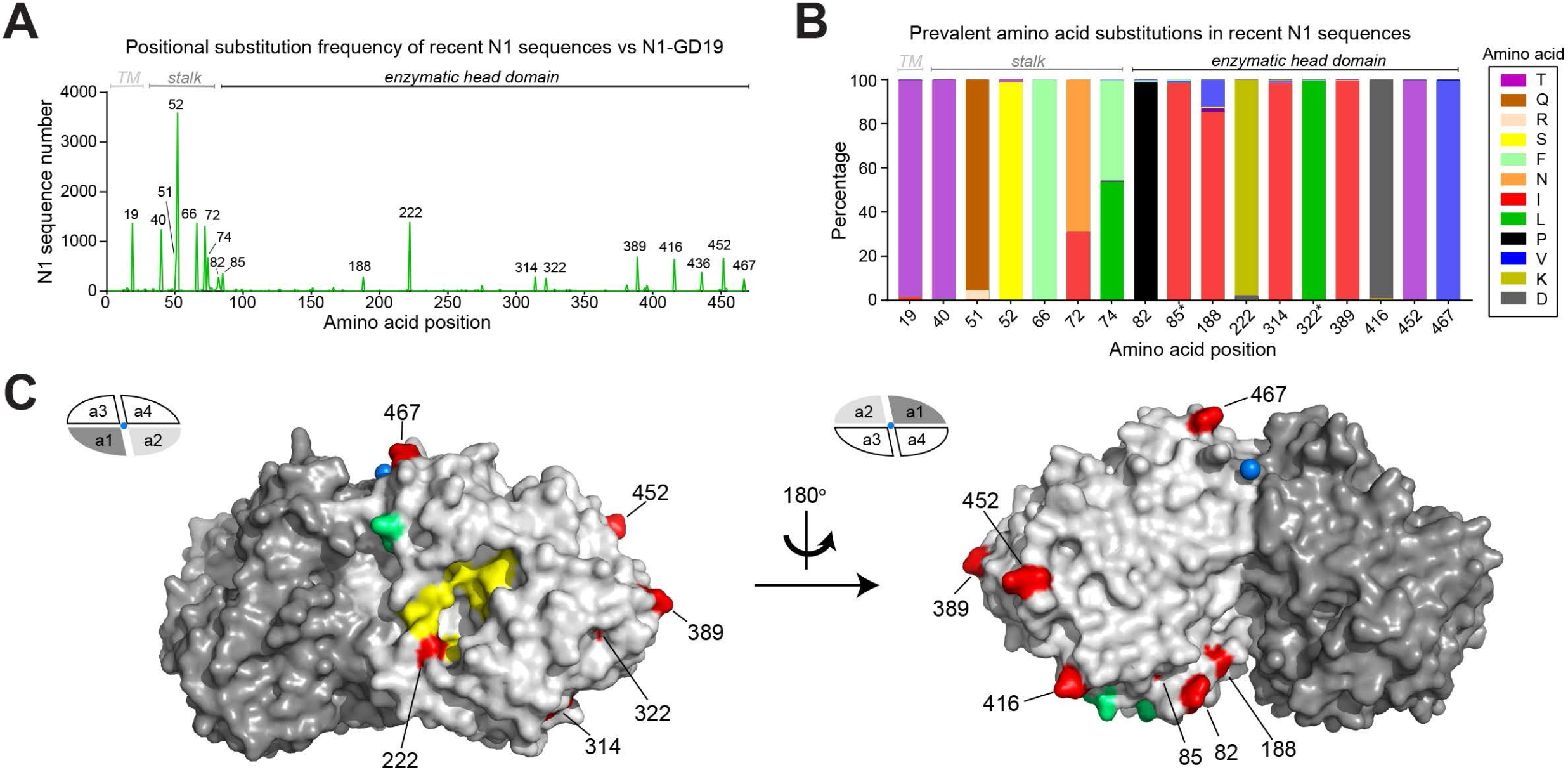
Positional variation in the NA sequences from recent human H1N1 viruses. **A.** Graph displaying the number of recent N1 sequences that carry different amino acids than N1-GD19 at each position. The N1 sequences were from human H1N1 IAVs collected between September 1, 2019 and December 15, 2020. Regions corresponding to the transmembrane (TM) domain, stalk and the enzymatic head are indicated. **B.** Bar graph displaying the prevalence of the amino acids in the recent N1 sequences that differ from N1-GD19 at each position. Only the amino acids that are visible in the graph are referenced in the legend. Asterisk indicates amino acids with minimal surface exposure. **C.** The amino acid positions that differ in the recent N1 sequences were highlighted (red) on a structure of the N1 head domain (PDB ID: 3NSS)[50] along with the Asn residues (green) of the conserved N-linked glycosylation sites [44], the NA active site residues (yellow), and the central Ca^2+^ ion (blue). The side view (left) of the NA tetramer and the oligomeric interface between NA dimers (right) were generated using Pymol.

Protective epitopes in NA have been almost exclusively mapped to the head domain [22, 23, 33, 40, 41]. Therefore, we determined the distribution of the recent N1 sequences with one or more amino acid changes in the head domain and identified the most prevalent position combinations that contained substitutions. The number of head domain substitutions in the recent N1 sequences ranged from 0 to 7 (Fig. 4A), and ~66% of the head domain sequences were either identical to N1-GD19, or possessed substitutions at 1, 2, 5, or 6 specific positions in the head domain (Fig. 4B). We then identified representative NAs for the four head domain substitution combinations (Fig. 4C), generated PR8 reassortant viruses, and examined the reactivity to ferret antisera against N1-GD19 and NAs from three earlier WHO recommended H1N1 IAV vaccine strains. The ELLA results indicated that the representative NAs, which are present in ~66% of circulating H1N1 strains, are antigenically similar to N1-GD19, as well as N1-BR18 and N1-MI15, but not N1-CA09 (Table 2). This suggests that N1 antigenicity has remained relatively stable since 2015 and that the N1 antigens in the cell and egg-based vaccines for the 2020-21 season are a reasonable choice.

**Figure 4.**
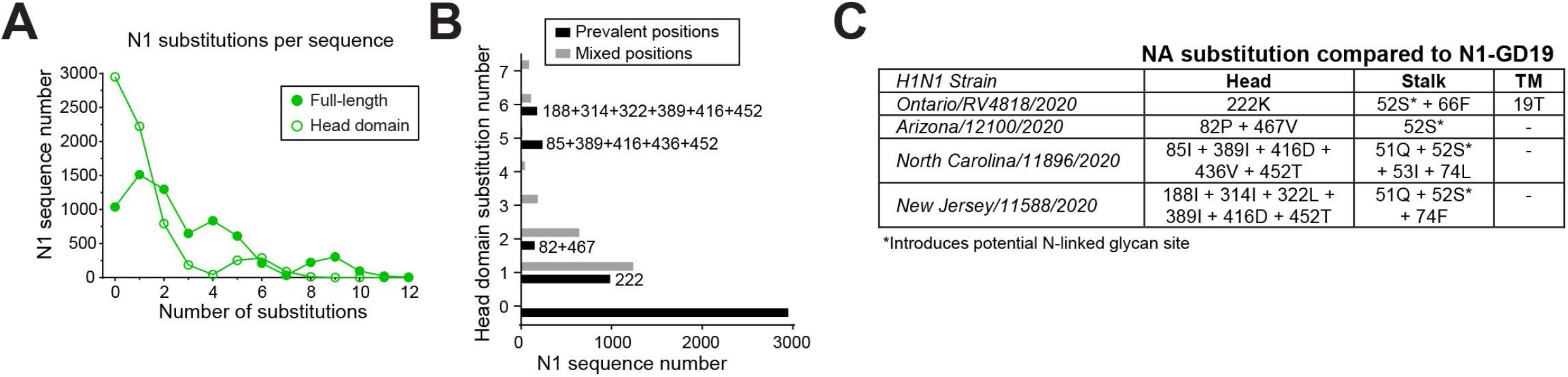
Sequence based selection of NAs from recent human H1N1 viruses. **A.** The number of N1 sequences from recent human H1N1 IAVs (isolated between 09-01-2019 and 12-15-2020) were plotted with respect to the number of amino acids that differ from N1-GD19. Distributions for both the full-length sequences and the sequences corresponding to the enzymatic head domain (residues 82-469) are displayed. **B.** N1 sequences were grouped based on the number of head domain substitutions with respect to N1-GD19. The N1 sequences from each group that possessed the indicated amino acid substitution sites in the head domain were extracted and tabulated (black) together with the number of sequences that possessed substitutions at other sites (grey). **C.** Chart containing the representative N1s that were selected for antigenic comparison based on the presence of the prevalent substitutions in the head domain. The amino acid substitutions with respect to N1-GD19 are listed by the domain in NA. TM - transmembrane domain.

**Table 2.** Reciprocal NAI titres for the indicated N1 reassortant viruses and antisera. NAI titres were determined by calculating the mean *IC_50_* of the indicated ferret antisera from duplicate ELLA runs. Cells where the NA in the PR8 reassortant virus matches the sequence of the NA used to produce the ferret antisera are highlighted. All ferret antisera were produced via intranasal inoculation with PR8 reassortant viruses carrying the indicated NA and H6.

|  |  | Ferret Antisera |  |  |  |
| --- | --- | --- | --- | --- | --- |
| | | α-N1-CA09 | α-N1-MI15 | α-N1-BR18 | α-N1-GD19 |
| Viruses | PR8<sup>N1-CA09</sup> | **798** | 70 | 110 | 256 |
|  | PR8<sup>N1-MI15</sup> | 193 | **785** | 1454 | 5376 |
|  | PR8<sup>N1-BR18</sup> | 142 | 534 | **1043** | 4750 |
|  | PR8<sup>N1-GD19</sup> | 149 | 582 | 1126 | **3884** |
|  | PR8<sup>N1-AZ20</sup> | 142 | 497 | 959 | 3478 |
|  | PR8<sup>N1-Ont20</sup> | 121 | 518 | 720 | 3082 |
|  | PR8<sup>N1-NC20</sup> | 141 | 469 | 871 | 3046 |
|  | PR8<sup>N1-NJ20</sup> | 95 | 271 | 548 | 1806 |

### Sequence and antigenic comparison of NAs from recent H3N2 and vaccine viruses

The positional analysis of the recent NA sequences from H3N2 IAVs was performed using both the N2-HK19e and N2-HK19c sequences as a reference (Fig. 5A). In contrast to N1, amino acid changes in the recent N2s were more common at positions in the head domain than in the stalk. We also observed that the C-terminal residue of N2-HK19e (469T) and the stalk residue of N2-HK19c (45S) are not common in the recent N2 sequences. The amino acid differences in all the prevalent stalk and head domain positions were mainly attributed to a specific amino acid, making it easier to determine if the substitutions influence N2 antigenicity (Fig. 5B). Most of the head domain substitutions mapped to surface exposed residues and many are positioned around the active site and on the top of the tetramer near the central Ca^2+^ binding site (Fig. 5C), which has been shown to be critical for NA activity [42].

**Figure 5.**
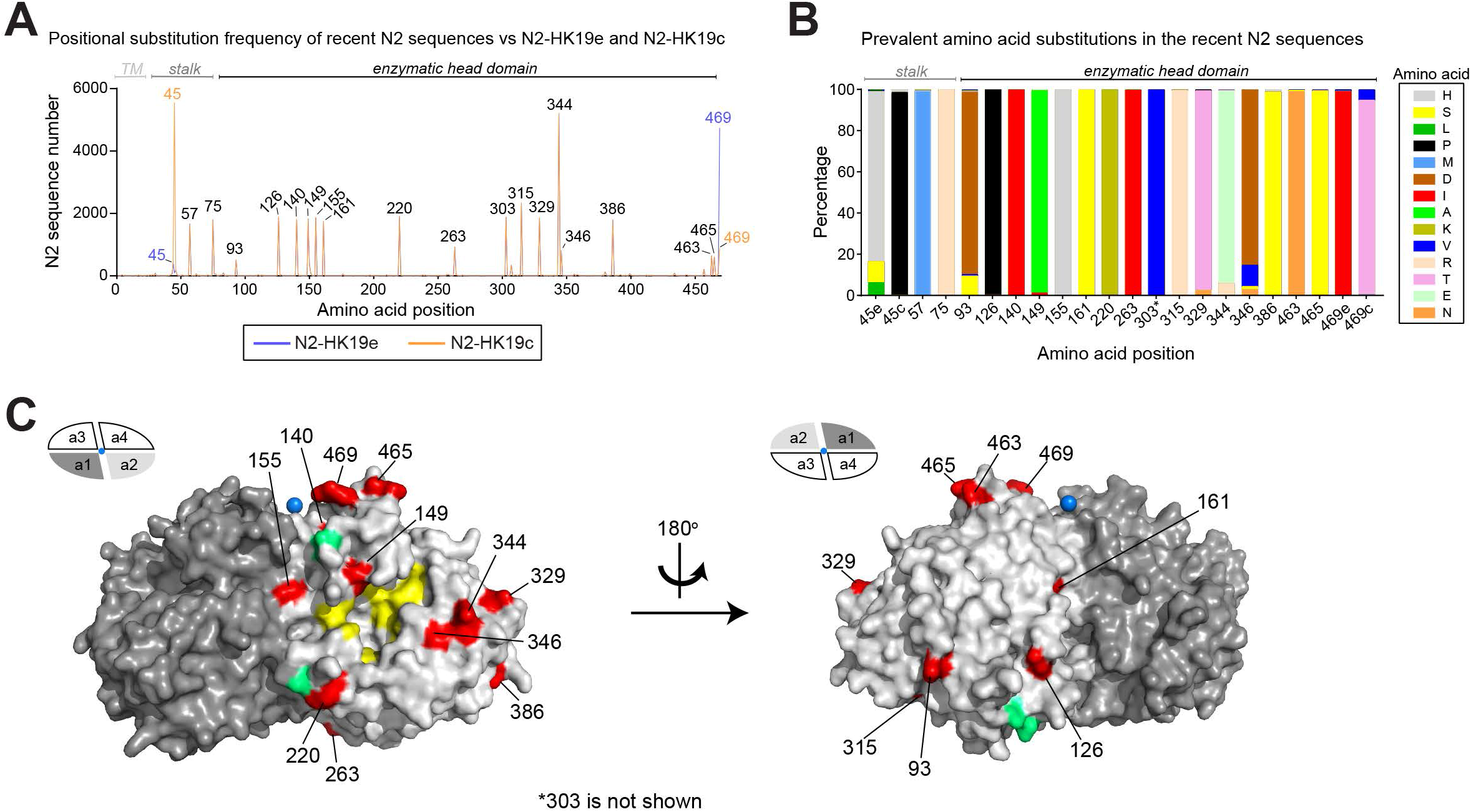
Positional variation in the NA sequences from recent human H3N2 viruses. **A.** Graph displaying the number of recent N2 sequences that carry different amino acids than N2-HK19c (orange) and N2-HK19e (blue) at each position. The N2 sequences were from the human H3N2 IAVs isolated between 09-01-2019 and 12-15-2020. Amino acid positions corresponding to the TM domain, stalk and the enzymatic head are indicated. **B.** Bar graph displaying the prevalence of the amino acids in the recent N2 sequences that vary from N2-HK19e and N2-HK19c at each position. Note N2-HK19e and N2-HK19c differ at position 45 and 469. Only amino acids visible in the graph are referenced. Asterisk indicates amino acids with little surface exposure. **C.** The amino acid positions that differ in the recent N2 sequences were highlighted (red) on a structure of the N2 head domain (PDB ID: 4GZT)[51] along with the Asn residues (green) of the conserved N-linked glycosylation sites, the NA active site residues (yellow), and the central Ca^2+^ ion (blue). The side view (left) of the NA tetramer and the oligomeric interface between NA dimers (right) were generated using Pymol.

Only a few of the recent N2 sequences were identical to N2-HK19e or N2-HK19c and the sequences formed a bimodal distribution based on the number of amino acid differences, which slightly shifted when the analysis was performed on the head domain sequences (Fig. 6A). Tabulating the most prevalent substitution combinations in the head domain (with respect to N2-HK19c) revealed that ~55% of the recent N2 sequences contain amino acid substitutions at 2, 4, 5, 6, or 10 specific positions in the head domain (Fig. 6B). PR8 reassortant viruses were generated that carry representative NAs with seven of the identified head domain substitution combinations (Fig. 6C) and the reactivity to ferret antisera against N2-HK19e, N2-HK19c, and the NAs from four earlier WHO recommended H3N2 IAV vaccine strains was examined.

**Figure 6.**
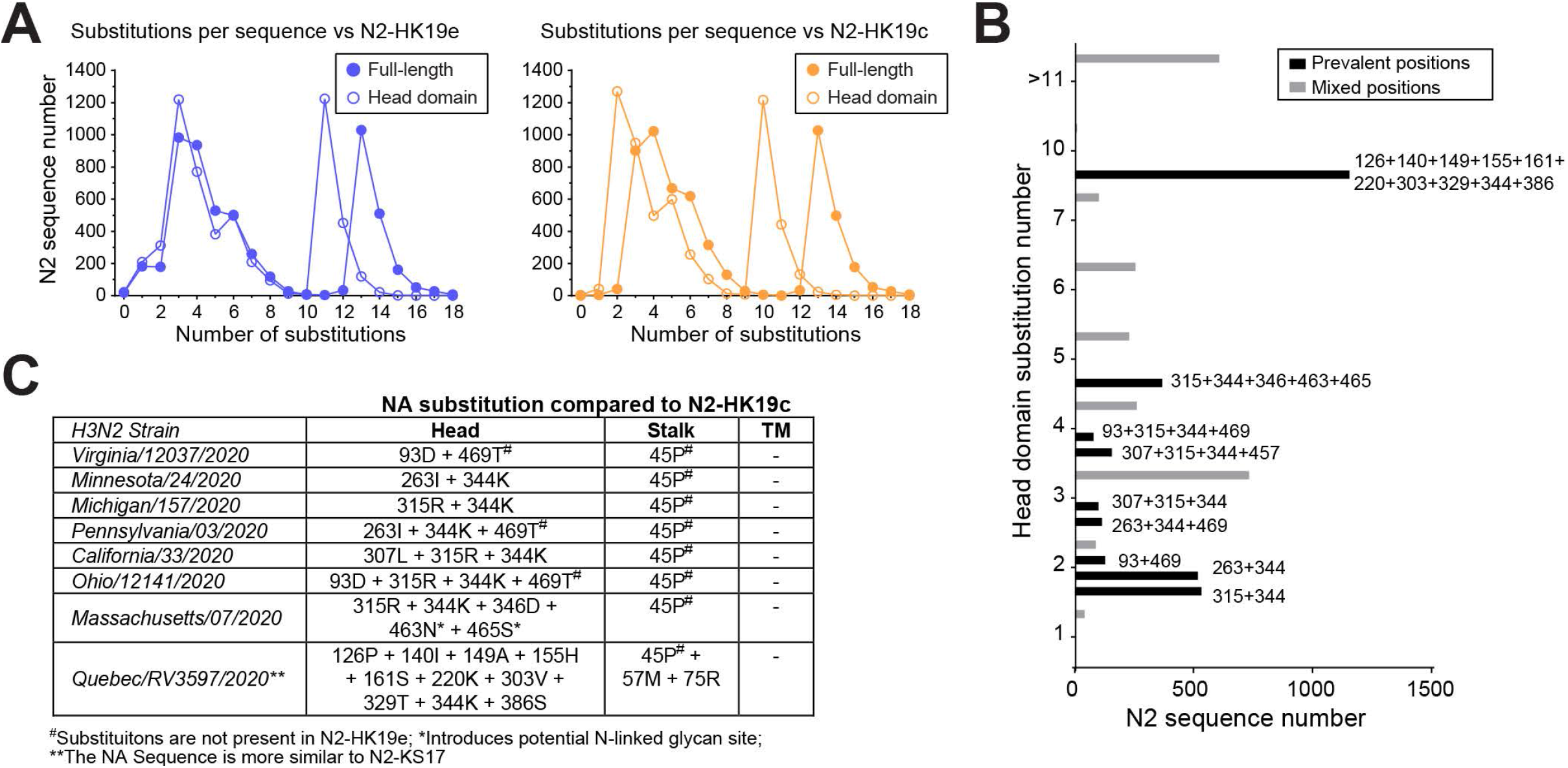
Sequence based selection of NAs from recent human H3N2 viruses. **A.** The number of N2 sequences from recent human H3N2 IAVs (isolated between September 1, 2019 and December 15, 2020) were plotted with respect to the number of amino acids that differ from N2-HK19e (left) and N2-HK19c (right). Distributions for both the full-length sequences and the sequences corresponding to the enzymatic head domain (residues 82-469). **B.** N2 sequences were grouped based on the number of head domain substitutions with respect to N2-HK19c. The N2 sequences from each group with the indicated amino acid substitution sites in the head domain (black) were extracted and tabulated together with the number of sequences that possessed substitutions at other sites (grey). **C.** Chart containing the representative N2s that were selected for further antigenic comparison based on the presence of the prevalent substitution in the head domain. The amino acid substitutions with respect to N2-HK19c are listed by the domain in NA.

The ELLA results indicated that the representative N2s, which are present in ~55% of circulating H3N2 strains, all possess reasonable antigenic similarity to N2-HK19e and N2-HK19c (Table 3). The high NAI titre for the N2 with 10 substitutions (N2-Que20) was not surprising as the sequence is almost identical to N2-KS17 and the antisera against N2-HK19e and N2-HK19c also generated very high NAI titres for PR8^N2-KS17^. Supporting that the C-terminus is part of an antigenic domain for NAI antibodies in N2, the N2-HK19e ferret serum continued to show higher reactivity than the N2-HK19c ferret serum for all the N2s possessing the 469T substitution (Table 3). Results from the other NA vaccine antisera suggest that the N2s since 2016 share some antigenic features, whereas those from 2013 and 2014 are more similar. Together, these data imply that the N2 antigens in the egg and cell-based vaccines for the 2020-21 season match reasonably well against the circulating H3N2 strains, which includes the large subpopulation of H3N2 viruses that carry a NA similar to N2-KS17. They also demonstrate that it may be more beneficial to choose H3N2 vaccine strains that possesses a NA with a common C-terminus.

**Table 3.**
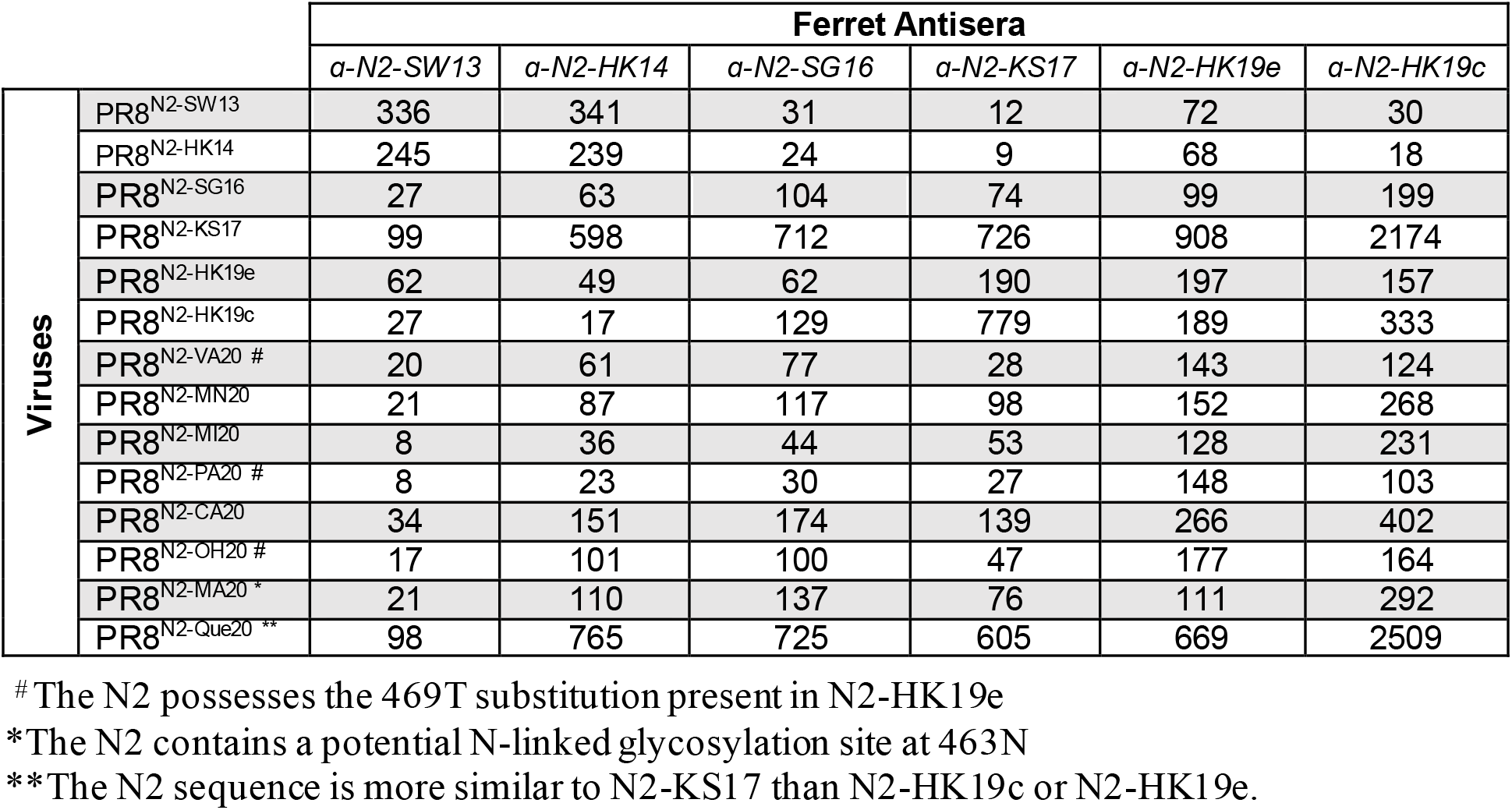
Reciprocal NAI titres from the indicated N2 reassortant viruses and antisera. NAI titres are the mean *IC_50_* values for the indicated ferret antisera from duplicate ELLA runs. Cells where the NA in the PR8 reassortant virus matches the sequence of the NA used to generate the ferret antisera are highlighted. All ferret antisera were produced by intranasal inoculation with PR8 reassortant viruses carrying the indicated NA and H6.

## DISCUSSION

During previous influenza seasons our lab has supported the influenza A vaccine strain selection process by using available isolates to evaluate the circulating NA antigens. The isolate dependence severely impaired these evaluations because only a few H1N1 and almost no H3N2 strains were received, partially due to the poor growth of recent H3N2 strains in eggs. Additionally, the few isolates that were obtained often arrived close to the February strain selection meeting and most carried identical NAs. To avoid these issues and perform a more systematic evaluation of the prevalent NAs in circulating strains, we changed our strategy for the 2020-21 season to a sequence-based approach that used reverse genetic viruses for the NA source. Our results showed that the N1 and N2 antigens in the recommended 2020-21 IAV strains for egg and cell-based vaccines are antigenically similar to most of the NAs in circulating H1N1 and H3N2 viruses, making them reasonable selections. In contrast to previous seasons, these conclusions were based on data from two panels that represented ~66% of the N1 and ~55% of the N2 sequences from the strains collected between September 2019 to December 2020, suggesting the results from this type of approach will likely provide insight on the majority of N1s and N2s that circulate during the remainder of an influenza season.

The data for the different N2s from the egg and the cell-based vaccine strains also revealed that the C-terminus of N2 is within or can influence an antigenic domain for NAI antibodies. This interpretation, which suggests the cell-based N2 with 469I could provide slightly better protection against most circulating strains, was based on the reproducible bias of the two antisera for N2s with an identical C-terminus and previous work demonstrating the amino acid at position 468 can influence NAI titres for N2 [43]. Interestingly, the N2 C-terminus is near the central Ca^2+^-binding site that was previously shown to be crucial for N1 enzymatic activity [42], raising the possibility that antibody binding decreases the Ca^2+^ affinity of this site, thereby reducing N2 activity. Although more directed studies are needed to determine the precise mechanism of inhibition, these observations indicate the N2 C-terminus should receive some consideration during the H3N2 strain selection. This especially applies to the 2021-22 season where the newly recommended H3N2 vaccine strain carries a NA with a C-terminal N-linked glycosylation site. The new glycosylation site at position 463 is distinct from the variable glycosylation sites that have previously been reported for N2 [44], and could impact any N2-mediated vaccine protection if the prevalent circulating N2s lack this site.

The only difference we observed for the recent N1s was the ~2-fold lower NAI titres for N1-NJ20 with all four antisera, suggesting substitutions at positions 188, 314, 322, 389, 416 and 452 may somewhat change N1 antigenicity. However, N1-NC20 carries three of the substitutions and showed no NAI titre decrease, implying that any potential antigenic change is caused by substitutions at positions 188, 314 or 322 alone, or in combination with the other three sites. While the potential selection of N1s with this constellation should receive some attention in the future, the antisera reactivity remained rather high indicating a N1-mediated response would still recognize this NA. It is also plausible that the use of two-fold serial dilutions in the ELLA contributed to the observed NAI titre variation.

An advantage of this approach is that relevant data and reagents were already available for evaluating the NAs from the new H1N1 and H3N2 vaccine strains that were recently recommended for the 2021-22 northern hemisphere influenza season. Specifically, the NA protein sequences from the recommended H1N1 strains for egg (A/Victoria/2570/2019) and cell-based (A/Wisconsin/588/2019) vaccines are both identical to N1-Ont20 that was tested. In addition, the NA sequence from the recommended H3N2 strain for egg and cell-based vaccines (A/Cambodia/e0826360/2020) only differs from the N2-MA20 that was tested by a single residue in the stalk (position 62), which is unlikely to change the antigenic properties. Another advantage is the accumulation of a thorough antigenic data set for NAs that have been prevalent in humans that can be utilized in the future to establish a comprehensive NA antigenic map which can help to improve the strain selection process.

There still are some significant deficiencies in the ELLA-based antigenic characterization of NA. The most striking are the false positive NAI titres that are obtained from antibodies against HA [38, 39], the time consuming nature of the assay, and the significantly lower NAI titres that are obtained for N2 compared to N1, resulting in a smaller dynamic range for evaluating N2. Despite these deficiencies, systematic approaches are likely the most effective way for evaluating antigenic drift in NA and the current ELLA has been shown to correlate reasonably well with protection in humans [21, 35], making it an applicable assay for examining NA antigenicity. Future modifications to alleviate one or more of these deficiencies will increase the likelihood that NA antigens in circulating viruses are evaluated as thoroughly as the HA antigens during the vaccine selection process.

## METHODS

### Reagents

Madin-Darby canine kidney 2 cells (MDCK.2; CRL-2936) and HEK 293T/17 cells (CRL-11268) were purchased from LGC Standards. Dulbecco’s Modified Eagles Medium (DMEM), fetal bovine serum (FBS), L-glutamine, penicillin/streptomycin (P/S), Opti-MEM I (OMEM), Simple Blue Stain, Novex 4-12% Tris-Glycine SDS-PAGE gels, Maxisorp 96-well plates and Lipofectamine 2000 transfection reagent were obtained from Thermo Fisher Scientific. 2’-(4-methylumbelliferyl)-α-d-*N*-acetylneuraminic acid (MUNANA) and Zanamivir were acquired from Cayman Chemicals and Moravek Inc. Bovine fetuin, *o*-phenylenediamine dihydrochloride (OPD), and HRP-linked peanut agglutinin were purchased from Sigma. Specific-pathogen-free (SPF) eggs and turkey red blood cells (TRBCs) were purchased from Charles River Labs and the Poultry Diagnostic and Research Center (Athens, GA), respectively.

### Plasmids and constructs

Reverse genetics (RG) plasmids containing the PR8 gene segments were provided by Dr. Robert Webster (St. Jude Children’s Research Hospital) have the following GenBank Identifications: CY038902.1 (PR8-PB2), CY038901.1 (PR8-PB1), CY084019.1 (PR8-PA), CY146825.1 (PR8-HA), CY038898.1 (PR8-NP), MH085246.1 (PR8-M), and CY038899.1 (PR8-NS). The RG plasmids containing N1-CA09, N1-MI15, N1-BR18 and the H6 gene (A/turkey/Mass/3740/1965) were described previously [42, 45, 46]. The genes corresponding the NAs from the following strains were synthesized and inserted into the pHW2000 plasmid [47] by Genscript: A/H1N1/Guangdong-Maonan/SWL1536/2019 (N1-GD19), A/H1N1/Ontario/RV4818/2020 (N1-Ont), A/H1N1/Arizona/12100/2020 (N1-AZ20), A/H1N1/North Carolina/RV4818/2020 (N1-NC20), A/H1N1/New Jersey/11588/2020 (N1-NJ20), A/H3N2/Hong Kong/45/2019 (N2-HK19c), A/H3N2/Hong Kong/2671/2019 (N2-HK19e), A/H3N2/Virginia/12037/2020 (N2-VA20), A/H3N2/Minnesota/24/2020 (N2-MN20), A/H3N2/Michigan/157/2020 (N2-MI20), A/H3N2/Pennsylvania/03/2020 (N2-PA20), A/H3N2/California/33/2020 (N2-CA20), A/H3N2/Ohio/12141/2020 (N2-OH20), A/H3N2/Massachusetts/07/2020 (N2-MA20), and A/H3N2/Quebec/RV3597/2020 (N2-Que20). All RG plasmids were confirmed by sequencing before use (FDA core facility).

### Cell culture, reverse genetics and viral propagation

MDCK.2 and HEK 293T/17 cells were maintained at 37 °C with 5% CO_2_ and ~95% humidity in DMEM containing 10% FBS and 100 U/ml P/S. Reassortant viruses were created by 8-plasmid reverse genetics [47] in 6-well plates using the indicated NA, or NA and HA pair, and the complimentary seven, or six, gene segments of PR8. The day before each well received ~1 × 10^6^ 293T and ~1 × 10^6^ MDCK.2 cells. The eight plasmids (1 μg of each) were added to 200 μl of OMEM, mixed with 18 μl of lipofectamine, and incubated 45 min at room temperature. The wells were washed with 2 ml OMEM, the mixture was added to each well and incubated 5 min at 37 °C prior to adding an additional 800 μl OMEM. An additional 1 ml OMEM containing 4 μg/ml TPCK trypsin was added to each well at 24 h post-transfection. Culture medium containing the rescued viruses was harvested ~96 h post-transfection and clarified by sedimentation (2000 × g; 5 min) prior to being used to inoculate 9-11 day old embryonic SPF chicken eggs (100 μl per egg). After incubating the eggs for 3 days at 33 °C they were placed at 4 °C for 2 h, the allantoic fluid was collected, clarified by centrifugation (2000 × g; 5 min), aliquoted and stored at −80 °C. Prior to use each viral stock was confirmed using MiSeq RNA deep sequencing (FDA core facility).

### Viral purification and SDS-PAGE

PR8^H1N1-BR18^ and WSN^H1N1-BR18^ reassortant viruses were propagated in SPF chicken eggs and purified as previously described [25]. The purified viruses (~5 μg total viral protein) were mixed with 2X sample buffer and heated at 50 °C for 5 min prior to being resolved on a 4-12 % polyacrylamide Tris-Glycine SDS-PAGE wedge gel. Gels were stained with simple blue and imaged with an Azure C600 Bioanalytical Imaging System (Azure Biosystems).

### Bio-layer interferometry, NA activity and HAU measurements

Biotinylated fetuin was made using the EZ-Link NHS-PEG4-Biotinylation kit (Thermo Scientific). Briefly, bovine fetuin (Sigma) was dissolved in PBS pH 7.2 to a concentration of 10 mg/ml and incubated with 2.9 mM of the biotinylation reagent at room temperature for 60 min. Biotinylated fetuin was then isolated using an Agilent 1260 Infinity II HPLC equipped with a 21.2 x 300 mm AdvanceBio SEC 300Å column, a variable wavelength detector and PBS 7.2 as the mobile phase. Fractions containing fetuin were pooled, the protein concentration was determined using BCA, and the labelling was calculated to be ~8 biotin/molecule of fetuin. Binding of the purified BPL inactivated PR8^H1N1-BR18^ and WSN^H1N1-BR18^ viruses to immobilized biotinylated fetuin was performed using a ForteBio Octet Red96 bio-layer interferometer equipped with ForteBio High Precision Streptavidin biosensors at 25 °C with a plate shake speed of 1000 rpm. Baselines were established for 2 min in 200 μl of buffer (PBS pH 7.2 containing 1mM CaCl_2_ and 0.005% Tween20) prior to and post incubation of the biosensor with 200 μl of 10 nM biotinylated fetuin for 10 min. Fetuin loaded biosensors were then transferred to wells containing 200 μl of the indicated concentrations of the WSN^H1N1-BR18^ or PR8^H1N1-BR18^ viruses in buffer with 10 μM zanamivir for 30 min to analyse HA-mediated virus association. The biosensors were then transferred to new wells containing 200 μl of buffer for 20 min to examine NA-mediate dissociation. NA activity and HAU measurements were performed using equal amounts of total viral protein for NA activity and HAU measurements as previously described [25].

### Ferret inoculation and serum collection

All animal experiments were approved by the U.S. FDA Institutional Animal Care and Use Committee (IACUC) under Protocol #2001-05. The animal care and use protocol meets National Institutes of Health (NIH) guidelines. Intranasal inoculations were performed on 14 week-old male ferrets using allantoic fluid of the indicated reassortant viruses. Approximately 0.5 ml of allantoic fluid was equally dispensed between both nostrils of the anesthetized ferret using a 1 ml syringe equipped with a 20 Gauge feeding needle. Ferret temperatures, monitored using an implanted microchip, and body weight were recorded daily for two weeks and the sera was harvested 3 weeks post-infection, aliquoted and stored at −80°C.

### Enzyme-linked lectin assay (ELLA) and NAI titre determination

The ELLA was performed in two parts on 96-well Maxisorp plates coated with 2.5 μg/well of bovine fetuin as previously described [31]. Briefly, the correct virus concentration for the ELLA was first determined using a 2-fold dilution series for each virus. The dilution series for each virus was added to the fetuin-coated plates and incubated at 37 °C overnight. Plates were washed and incubated with ~100 ng/well of horse radish peroxidase-linked peanut agglutinin (PAG-HRP) for 2 h at room temperature. Plates were washed again and developed with OPD (0.5 ng/well) for 10 min, the reaction was stopped with 100 μl/well 1N sulfuric acid and the absorbance at 490 nm was measured. The absorbance was plotted versus the virus dilution and the dilution corresponding to the top of the linear part of the titration curve was chosen. Next, a 2-fold dilution series was made for each antiserum and transferred to a fetuin-coated plate prior to adding the virus at the concentration determined in the first assay. The plates were incubated at 37 °C overnight, washed, incubated with PAG-HRP and developed as described above. The absorbance for each antiserum and virus pair was background corrected using the average signal obtained from wells containing no virus or serum. To calculate the percent NA inhibition each background corrected sample signal was subtracted from the average virus control well signal (wells that received virus but no antiserum) and the difference was divided by the average virus control well signal and multiplied by 100%. The NA inhibition percentage was then plotted by the reciprocal dilution of the antiserum in GraphPad Prism 8.0 and the IC_50_ value, which corresponds to the NAI titre, was determined by a four-parameter least squares fit analysis.

### Sequence analysis

The available NA protein sequences collected from human H1N1 (N=6844) and H3N2 (N=5566) viruses since September 1, 2019 were downloaded from the Global Initiative on Sharing All Influenza Data (GISAD) database on December 15, 2020. Sequences longer or shorter than 469 amino acids were excluded along with those that contained ambiguous amino acid identities. All analyses were performed based on positions in the N1 or N2 amino acid sequences using the indicated vaccine NA sequence as a reference. Results specific for the head domain were obtained by examining the residues corresponding to amino acids 82 to 469, excluding the stalk and transmembrane domain [48, 49].

### Statistical analysis

*P* values were calculated with GraphPad Prism 8 software using Student’s unpaired t-test with a two-tailed analysis and a confidence interval (CI) of 95%. *P* values that were above 0.05 (not significant) were not included in any figures.

## DATA AVAILABAILITY

Data generated for this study that support the reported findings are included in the published article. Additional supporting data are available from the corresponding author.

## ACKNOWLEDGEMENTS

We would like to thank the Global Initiative on Sharing All Influenza Data (GISAID) for providing the sequence data necessary for this evaluation and members of the Division of Viral Products at the Center for Biologics Evaluation and Research (CBER), as well as Dr. Maryna Eichelberger (FDA) for the critical reading of the manuscript and offering several helpful suggestions. This work was supported by Intramural funding to the lab from CBER at the US Food and Drug Administration (FDA).

